# BmHen1 plays an essential role in the regulation of eupyrene sperm development in *Bombyx mori*

**DOI:** 10.1101/2022.06.30.498356

**Authors:** Xu Yang, Dongbin Chen, Shirui Zheng, Meiyan Yi, Zulian Liu, Yongjian Liu, Dehong Yang, Yujia Liu, Linmeng Tang, Chenxu Zhu, Yongping Huang

## Abstract

In lepidopteran insects, sperm polymorphism is a remarkable feature, in which males exhibit two different types of sperms. Both sperm morphs are essential for fertilization as eupyrene (nucleate) sperm carries DNA and fertilizes the egg, while apyrene (anucleate) sperm is necessary for transporting eupyrene sperm into females. To date, the functional genetic study on dichotomous spermatogenesis has been limited. It is known that, in the model species including mice, worms, and flies, the components in piRNA biogenesis pathway play an important role in gonad development. In this study, we characterize BmHen1 as a new critical component involved in the regulation of eupyrene sperm development in *B. mori*. We generated the loss-of-function mutant of *BmHen1* (*ΔBmHen1*) through CRISPR/Cas9-based gene editing, and found that it is both female- and male-sterile. *ΔBmHen1* females lay significantly fewer eggs than wild-type, which display morphological defects. Fluorescence staining assays show that the *ΔBmHen1* eupyrene sperms exhibit severe defects in nuclei formation, while its apyrene sperms are normal. We then constructed the loss-of-function mutants of *Siwi* and *BmAgo3* (*ΔSiwi* and *ΔBmAgo3*) through CRISPR/Cas9-based gene editing, which encode PIWI proteins acting as the core elements in piRNA biogenesis, and explored whether they might be involved in spermatogenesis. To our surprise, *ΔSiwi* and *ΔBmAgo3* mutants develop normal male reproduction system, indicating that they don’t participate in sperm development. As the activity of BmHen1 depends on BmPnldc1 during piRNA biogenesis, and *ΔBmHen1* and *ΔBmPnldc1* mutants display similar defects in sperm development, we performed RNA sequencing analysis to look for the genes that might be co-regulated by BmHen1 and BmPnldc1. Our results indicate that the defects in *ΔBmHen1* and *ΔBmPnldc1* eupyrene sperms could be attributed to dysregulated genes involved in energy metabolism and cell differentiation. Furthermore, we found that the piRNA biogenesis is inhibited in *ΔBmHen1* and *ΔBmPnldc1* sperm bundles, whereas the transposon activity was induced. Taken together, our findings suggest that BmHen1 is a new crucial component regulating eupyrene sperm development in *B. mori*, whereas the PIWI proteins Siwi and BmAgo3 are not involved in this process. Our results may provide a potential gene target for genetic modification of sterility in *B. mori*.

## 1. Introduction

Spermatogenesis is a fundamental biological process for sexual reproduction species (Lesch and Page, 2012). During this process, germ cells undergo mitosis and meiosis, and complete morphogenesis changes to develop into sperms (Fuller, 1998; Kanippayoor et al., 2013). Sperms exhibit a remarkable diversity in morphology and molecular levels (Dallai et al., 2016; Dumser, 1980; Lie et al., 2009). In Lepidoptera, sperm polymorphism is a fascinating feature, in which two morphs of sperms, eupyrene sperm (nucleate) and apyrene sperm (anucleate), exist in one single male (Friedländer, 1997; M.Phillips, 1971). The eupyrene sperm carries DNA and fertilizes the egg, while the apyrene sperm assists the eupyrene sperm migration (S. Chen et al., 2020; Sakai et al., 2019). The studies on dichotomous spermatogenesis have predominantly been limited to the aspects related to cytology and developmental timing (Friedländer and Wahrman, 1971; Kawamura et al., 1998; Yamashiki and Kawamara, 1998, 1997). As genome editing has currently been feasible in the Lepidoptera insect, *Bombyx mori* (*B. mori*), it becomes a model insect to investigate spermatogenesis (Wang et al., 2013). However, up to date, only few genes including *poly(A)-specific ribonuclease-like domain-containing 1* (*BmPnldc1*), *Sex-lethal* (*BmSxl*), and *Maelstrom* (*BmMael*) have been experimentally linked to dichotomous spermatogenesis (Chen et al., 2019; S. Chen et al., 2020; Sakai et al., 2019). Therefore, the molecular mechanism underlying spermatogenesis is not well understood in *B. mori*.

PIWI-interacting RNAs (piRNAs) are a class of small RNAs of 24-31 nucleotides in size produced from transposons and discrete genomic loci called piRNA clusters (Ozata et al., 2019). The piRNAs guide a subset of Argonaute proteins with endonucleolytic activity named PIWI proteins (PIWIs) including Piwi, Aub, and Ago3 in *Drosophila* to target transcripts (Czech et al., 2018; Iwasaki et al., 2015; Kirino et al., 2010). PIWIs exist across the animal kingdom and display differences in copy numbers and structures (Iwasaki et al., 2015). piRNAs and PIWIs mainly function in germ cells during gametogenesis to suppress transposable elements, the selfish genomic elements that jump around the genome at the transcription level (Iwasaki et al., 2015). Over the past two decades, the components participating in piRNA biogenesis such as Gtsf1, Mael, Papi, Tud, Pnldc1, and Hen1 have been identified in model species including mice, worms, and flies (Czech et al., 2018). Mutations in piRNA biogenesis pathway typically result in sterility with stereotypical phenotypes in the female or male germline (Juliano et al., 2011). In *B. mori*, the biochemical functions of the components involved in piRNA biogenesis such as PIWIs (Siwi and BmAgo3), and BmPnldc1, BmHen1 and BmPapi are well studied in ovary-derived cells (BmN) (Sakakibara and Siomi, 2018), and some of them have been physiologically characterized in vivo (Chen et al., 2019; K. Chen et al., 2020; S. Chen et al., 2020; Li et al., 2018; Yang et al., 2021). For example, BmMael is involved in the regulation of both morphs of sperm development, while BmPnldc1 is required for eupyrene sperm development.

Hua Enhancer 1 (Hen1) was first identified in the model plant *Arabidopsis*, which is methyltransferase that modifies 2’-O-methylation at the 3’ end of small RNAs (Yu et al., 2005). Hen1 is then found in animals where it largely modifies the 2’-O-methylation at the 3’ end of piRNAs rather than that of small RNAs (Horwich et al., 2007; Kirino and Mourelatos, 2007; Modepalli et al., 2018; Peng et al., 2018; Saito et al., 2007). As an exception, the *Drosophila* Hen1 homolog modifies both siRNAs and piRNAs (Horwich et al., 2007; Saito et al., 2007). Hen1-mediated modification of piRNAs protects them from being digested by the endonuclease, thus promoting their stability and activating transposon activity (Izumi et al., 2016; Wang et al., 2016). Although Hen1 is expressed predominantly in gonads, its physiological functions differ among species. For example, *Drosophila* Hen1 is not involved in female or male fertility (Wang et al., 2016), while Zebrafish Hen1 is required for female germline development only (Kamminga et al., 2010). Whether BmHen1 plays a role in spermatogenesis in *B. mori* remains unknown.

In this study, we first characterized the function of BmHen1 in regulating spermatogenesis in *B. mori* by generating the loss-of-function mutants of *BmHen1* (*ΔBmHen1*) through CRISPR/Cas9-based gene editing. We found that *ΔBmHen1* mutant is both female- and male-sterile. *ΔBmHen1* females produce much less eggs than wild-type and show a misshapen small phenotype. Fluorescence staining assays demonstrate that *ΔBmHen1* males have severe defects in their nuclei, which are dislocalizated in the eupyrene sperm bundles. We further found that *ΔBmHen1* males show defects in eupyrene sperm bundles at the early elongating stage. We further constructed the loss-of-function mutants of *Siwi* and *BmAgo3* through CRISPR/Cas9-based gene editing, and found that their males are fertile, and that both morphs of their sperm bundles are normal. RNA-seq analysis of *ΔBmHen1* and *ΔBmPnldc1* mutants indicates that the defects in their eupyrene sperms could be attributed to dysregulated genes involved in energy metabolism and cell differentiation. Taken together, our results demonstrate that BmHen1 is essential for eupyrene sperm development in *B. mori*, whereas PIWI proteins are not.

## 2. Materials and methods

### 2.1 Silkworm strains

The multivoltine, nondiapausing *B. mori* strain, Nistari, was used as a wild-type (WT) in this study. Larvae were reared on fresh mulberry leaves under standard conditions at 25□. The *ΔBmPnldc1* mutant was generated previously (Chen et al., 2020).

### 2.2 RNA isolation, cDNA synthesis and Quantitative real-time PCR (qRT-PCR)

Total RNA was extracted from three individual mutants or WT animals using the TRIzol reagent according to the manufacturer’s instructions (YEASEN). An aliquot of 1 μg of the total RNA was used to synthesize cDNA using PrimeScript™ RT reagent Kit with gDNA eraser (Takara). qRT-PCR analysis was performed on a StepOnePlus Real-Time PCR system (Applied Biosystems) with a SYBR green Real-Time PCR master mix (Toyobo). The *B. mori ribosomal protein 49 gene* (*Bmrp49*) was used as an internal control to standardize the variation of different templates. The amplification program was as follows: the samples were incubated at 95□ for 5 min, followed by 40 cycles of 95□ for 15 s, and 60□ for 1 min. Sequences of the qRT-PCR primers are listed in supporting information Table 1.

**Table 1.**
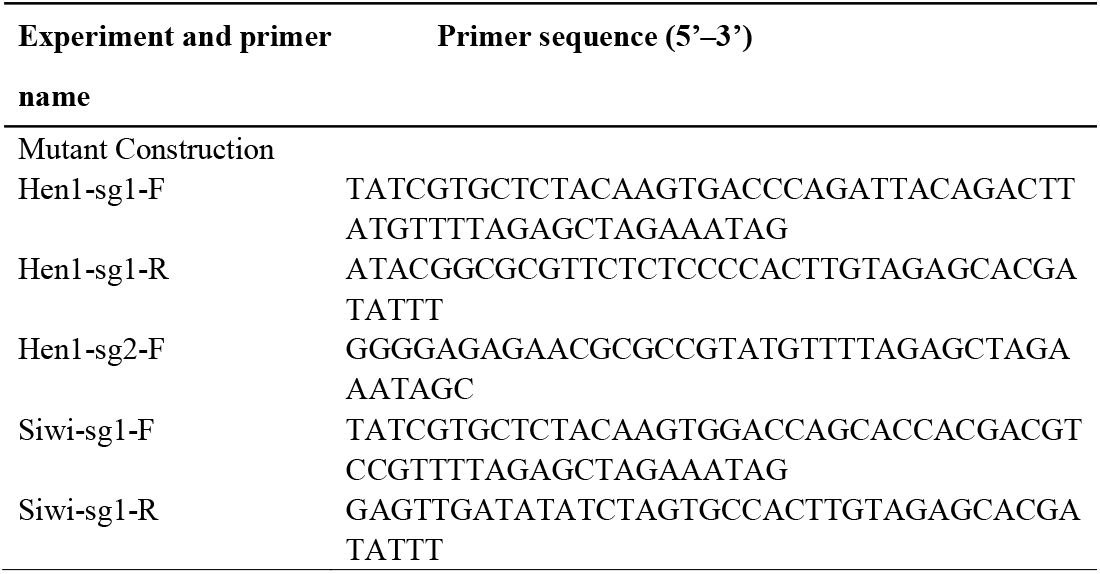
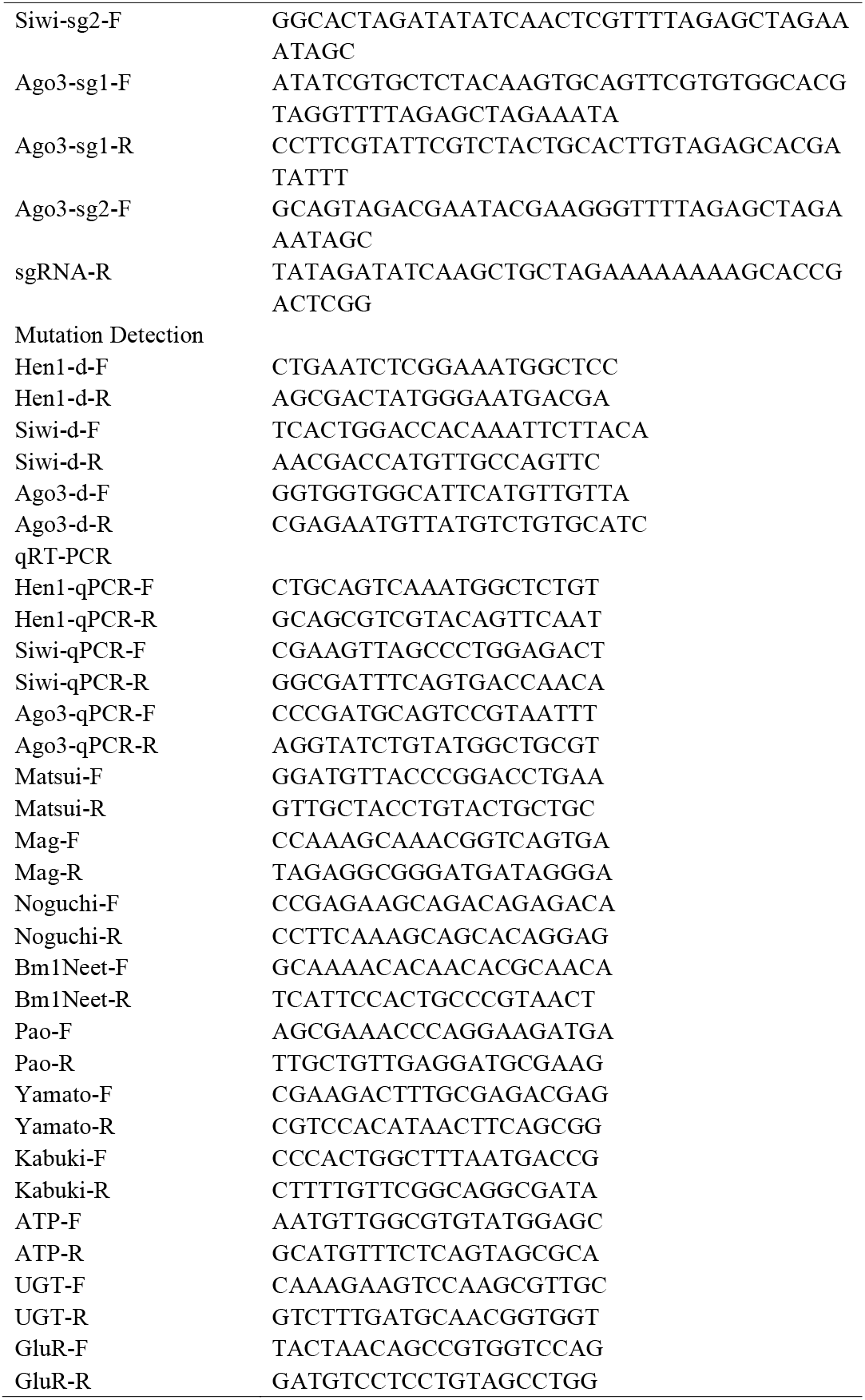
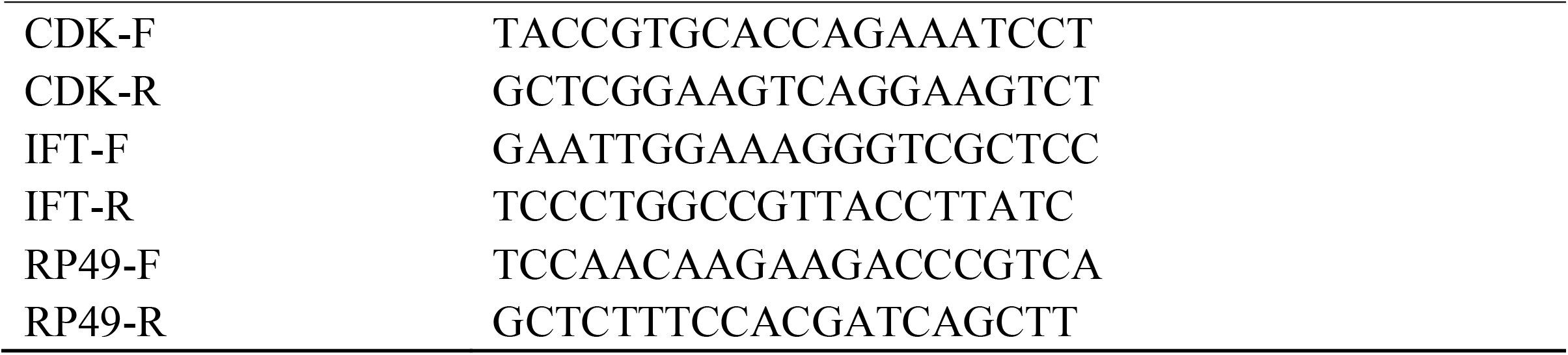
Primers used in this study

### 2.3 B. mori germline transformation and CRISPR/Cas9-mediated construction of BmHen1, Siwi and BmAgo3

A binary transgenic CRISPR/Cas9 system was used to construct the loss-of-function mutants of *BmHen1, BmSiwi*, and *BmAgo3* (*ΔBmHen1, ΔBmSiwi*, and *ΔBmAgo3*). The nos-Cas9 transgenic *B. mori* lines (nos-Cas9/IE1-EGFP) which express the Cas9 nuclease under the control of the *B. mori nanos* promoter was reported previously (Xu et al., 2017). The plasmids for expression of the two sgRNAs targeting each gene (U6-sgRNA /IE1-DsRed) under the control of the U6 promoter were constructed to generate *ΔBmHen1, ΔSiwi*, and *ΔBmAgo3*. Primers for plasmid construction and sgRNA targeting sequences are listed in Table 1.

For *B. mori* germline transformation, preblastodermal embryos were prepared and microinjected with transgenic plasmids (400 ng/μl) together with helper plasmids (IFP2, 200 ng/μl), and then incubated in a humidified chamber at 25□ for 10-12 days until hatching (Tamura et al., 2000; Tan et al., 2013). Larvae were reared with fresh mulberry leaves under standard conditions. Putative transgenic generation 0 (G0) moths were sib-mated or backcrossed with WT moths and G1 progenies were screened during early larval stages according to the GFP fluorescent marker visualized with a fluorescence microscopy (Nikon AZ100, Japan).

The nos-Cas9 lines and the U6-sgRNA lines were crossed to generate *ΔBmHen1, ΔBmSiwi*, and *ΔBmAgo3* with both EGFP and DsRed fluorescence markers for the subsequent experiments. Genomic DNA of the mutated animals was extracted by standard SDS lysis-phenol treatment, incubated with proteinase K, and purified for mutagenesis analysis via PCR amplification with specific primers (Table 1).

### 2.4 Fluorescence staining of sperm bundles

Immunostaining staining experiments were performed using spermatocysts and sperm bundles isolated from excised testes (fifth instar larvae stage to adult stage). The collected sperms were fixed for 1 h (Beyotime). After washed two times with PBS buffer, the samples were incubated with TRITC Phalloidin (YEASEN) for 1 h and then with Hoechst (Beyotime) for 20 min at room temperature. These samples were further washed three times with PBS buffer, and subsequently mounted in the antifade medium (YEASEN). All images were taken with an Olympus FV1000 microscope.

### 2.5 RNA-sequencing (RNA-seq) analysis

Total RNA from the sperm bundles on day four at the fifth larval stage was extracted from six individual animals of WT, *ΔBmHen1*, and *ΔBmPnldc1* and mixed together, respectively. The sperm bundles were released by tearing testes in PBS buffer. For mRNA sequencing, the total RNA that was firstly enriched and then fragmented was used for cDNA synthesis and library construction. The library was sequenced using Illumina 2000 platform, and the raw data were qualified, filtered, mapped, and quantified (FastQC, Trimmomatic, Bowtie2, Rsem) to the reference *B. mori* genome database (http://silkbase.ab.a.u-tokyo.ac.jp/cgi-bin/index.cgi) (Bolger et al., 2014; Kawamoto et al., 2019; Langmead and Salzberg, 2012; Li and Dewey, 2011). mRNA abundance was normalized with Deseq2. The calculated gene expression levels were then used to compare gene expression differences among samples (Y/X). Differentially expressed genes (DEGs) were screened based on the Poisson Distribution Method with a false discovery rate (FDR) < 0.05 and the absolute value of log_2_(Y/X) > 1. Enrichment analyses of DEGs were conducted using the Kyoto Encyclopedia of Genes and Genomes (KEGG) database (org.bmor.eg.db). The visualization was processed by using R packages (ggplot2, pheatmap). For small RNA sequencing, RNAs ranging in size between 15-50 nt were gel-purified and used for library construction and sequencing. The generated reads were filtered and mapped to the piRbase (http://bigdata.ibp.ac.cn/piRBase/index.php) (Wang et al., 2019). The distribution length of the piRNA was analyzed by using Perl scripts.

### 2.6 Statistical analysis

All experiments in this study were performed with at least three replicates (except the RNA-seq). All the data were expressed as the mean ± standard error of the mean (SEM). Differences between groups were examined using either two-tailed Student’s t-test or two-way analysis of variance. All statistical calculations and graphs were made with GraphPad Prism version 9.

## 3. Results

### 3.1 Construction of loss-of-function mutants of BmHen1 through a binary CRISPR/Cas9 system

To explore the biological function of *BmHen1*, we used a binary transgenic CRISPR/Cas9 system to obtain its loss-of-function mutant (Fig 1A), *ΔBmHen1*. To this end, we designed two small guide RNAs (sgRNAs) targeting exons 3 and 6 of *BmHen1* (Fig 1B), and *ΔBmHen1* mutants were generated through genetic crossing between the U6-sgRNA lines and the nos-Cas9 lines (Fig 1A). Mutations in the randomly selected representative F1 offspring were detected by genomic PCR and DNA sequencing, which indicates that the ORF of *BmHen1* was shifted in both female and male *ΔBmHen1* individuals (Fig 1C). Further qRT-PCR analysis showed that *BmHen1* was hardly expressed in *ΔBmHen1* (Fig 1D and E). These results demonstrate that we successfully obtained the loss-of-function mutant of *BmHen1*.

**Fig 1.**
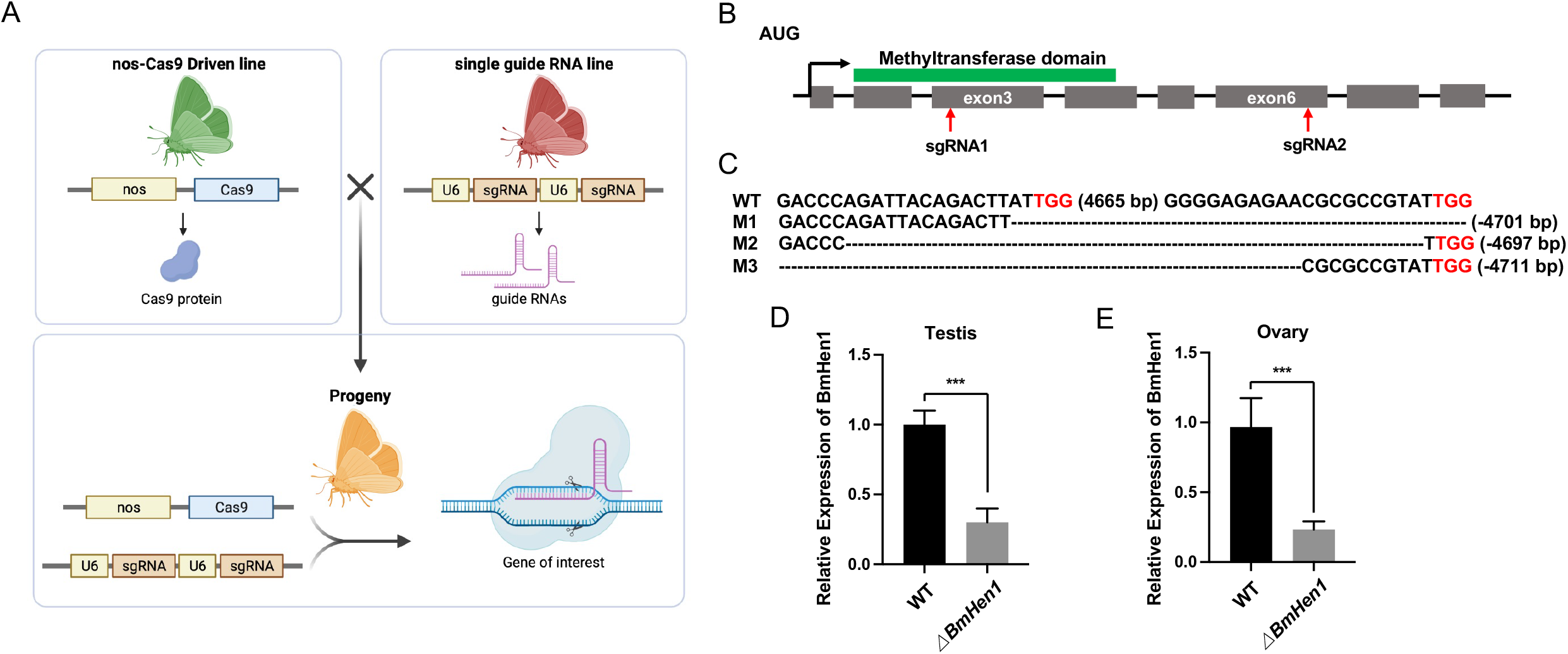
Construction of loss-of-function mutants of *BmHen1* using binary transgenic CRISPR/Cas9. (A) Schematic representation of mutant construction by using the binary transgenic CRISPR/Cas9 system. nos-Cas9 (green) transgenic moths were crossed with U6-sgRNA (red) transgenic moths and the F0 heterozygotes (yellow) were analyzed. (B) Genomic disruption of the BmHen1 gene using CRISPR/Cas9. Schematic of *BmHen1* gene structure and sgRNA targets. Exons of the ORFs are represented by gray filled boxes, and the function domain predicted by SMART is highlighted with a green box above. Red arrows indicate the target sites of sgRNA1 and sgRNA2. (C) Genomic mutations of the *BmHen1* gene are shown with target sequences. The dashed lines indicate deleted sequences. M1-M3 represents three independent mutant lines. (D) and (E) qRT-PCR analysis of the transcript levels of *BmHen1* in testis and ovary of WT and the loss-of-function mutant of *BmHen1* (*ΔBmHen1*) at the wandering stage. Three individual biological replicates were performed, and error bars are mean ± SEM. The asterisks (***) indicate significant differences relative to WT (*P* < 0.001, t-test).

### 3.2 Mutation of BmHen1 leads to both female and male sterility in Bombyx mori

The *ΔBmHen1* adults were viable and grossly normal, but the *ΔBmHen1* females laid much fewer eggs than the wild-type (WT) (Fig 2A and C). We found that *ΔBmHen1* mutant displayed morphological changes in the oviducts, some of which had no eggs or smaller or irregular eggs (Fig 2B). We then performed fecundity tests, and found that the crossing between *ΔBmHen1* female and WT male or between *ΔBmHen1* male and WT female generated the hybrids showing both female and male sterility (Fig 2D). These results demonstrate that *BmHen1* is essential for both female and male fertility.

**Fig 2.**
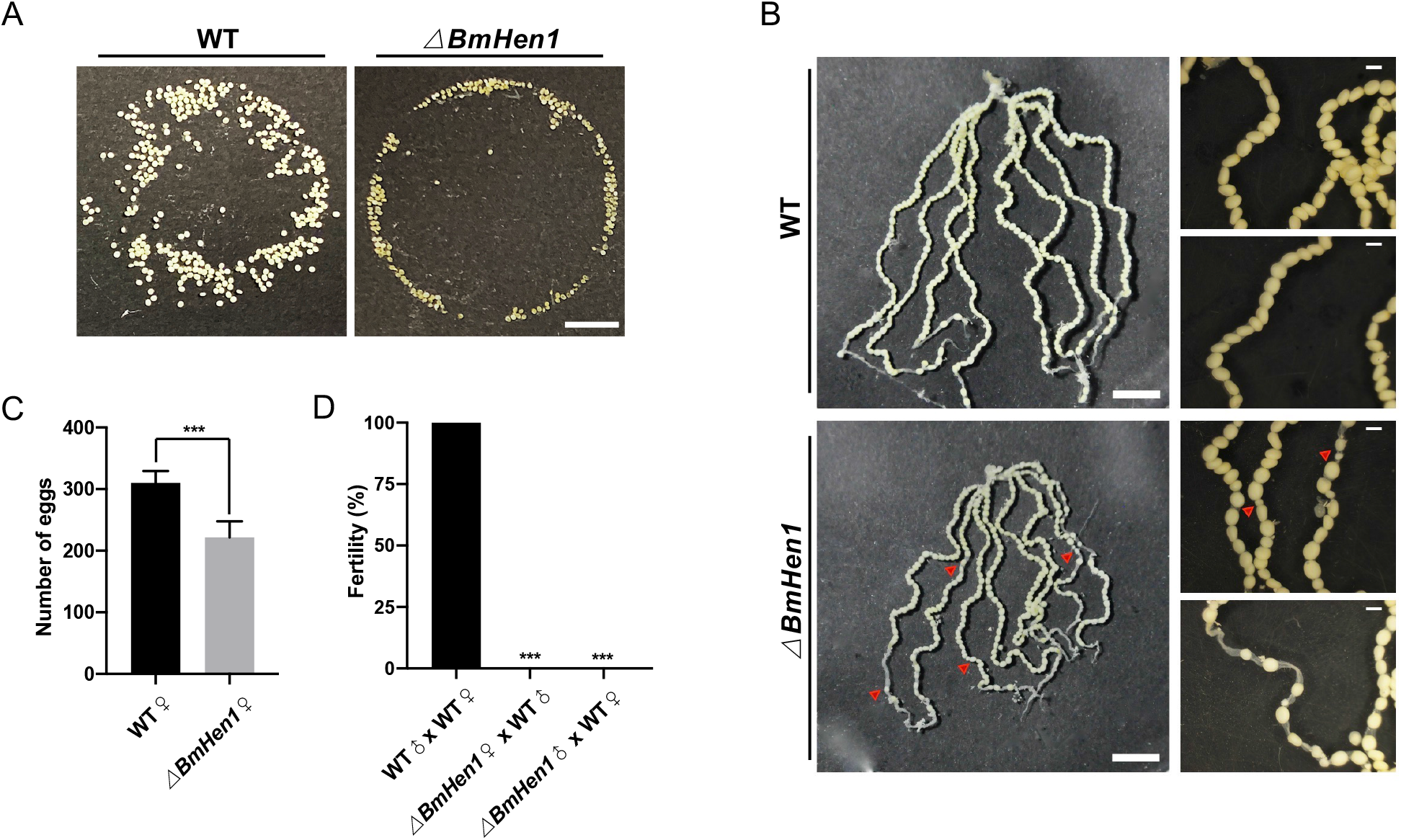
Mutation of *BmHen1* leads both male and female sterility. (A) Images of eggs laid by WT and *ΔBmHen1*. (B) Morphological analysis of oviducts of WT and *ΔBmHen1* moths. (C) Quantification of eggs laid by WT and *ΔBmHen1* (n=6, *P* < 0.001, t-test). (D) Fertility of males and females of the indicated genotypes. Fertility is indicated on the histogram (n= 15, *P* < 0.001, Fisher exact test). Scale bar, 1 cm.

### 3.3 BmHen1 is essential for eupyrene sperm development

It is known that the eupyrene and apyrene sperms of *B. mori* show different morphology and timing of differentiation during spermatogenesis (Kawamura et al., 1998; Yamashiki and Kawamara, 1998, 1997). Specifically, the eupyrene spermatogenesis begins on the first two days at the fifth instar larval stage, while the apyrene spermatogenesis starts during the wandering stage. As the male sterility observed for *ΔBmHen1* might result from the defects in the reproduction system (Fig 2D), we explored whether BmHen1 would be required for spermatogenesis. To do this, we first performed fluorescence staining to examine the development of both eupyrene and apyrene sperm bundles on the seventh day at the pupal stage in WT and *ΔBmHen1*. The results showed that the needle-shaped sperm nuclei were assembled regularly at the head of the eupyrene sperm bundles in the WT, whereas *ΔBmHen1* eupyrene sperm nuclei were abnormally organized and exhibited squeezed eupyrene sperm bundles (Fig 3A). No differences were detected in the apyrene sperm between WT and *ΔBmHen1* (Fig 3B).

**Fig 3.**
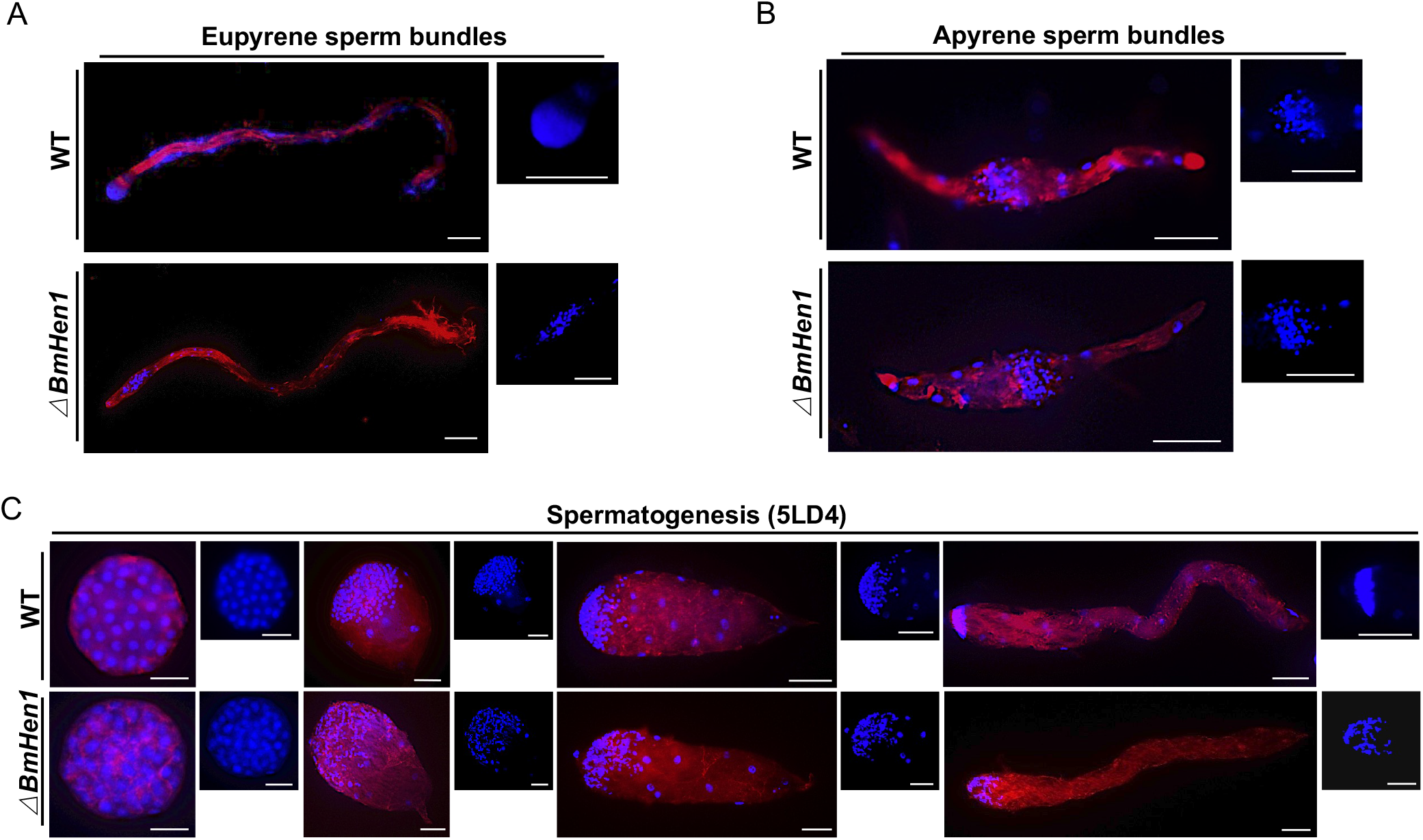
Mutation of *BmHen1* results in eupyrene sperm defects. (A and B) Representative immunostaining images of eupyrene sperm bundles (A) and apyrene sperm bundles (B) from WT and *ΔBmHen1* on the seventh day of the pupal stage. Scale bars, 50 μm. (C) Representative immunostaining images of eupyrene sperm bundles at different developmental stages on day four of the fifth larval instar of WT and *ΔBmHen1*. Scale bars, 50 μm. Blue, Hoechst; red, F-actin.

Next, we investigated eupyrene spermatogenesis in the testis on the fourth day of the fifth larval instar in WT and *ΔBmHen1* in details. The results showed that, at the early elongating stage, *ΔBmHen1* and WT developed similar eupyrene sperm bundles, with round nuclei being localized at the anterior part of the bundles (Fig 2C). At the late elongating stages, however, the eupyrene sperm nuclei were severely dislocated in *ΔBmHen1*, but not in WT (Fig 3C). Taken together, these results demonstrate that BmHen1 plays a vital role in the regulation of eupyrene spermatogenesis.

### 3.4 PIWIs proteins Siwi and BmAgo3 are dispensable for spermatogenesis in Bombyx mori

As PIWI proteins act as the core components in the piRNA biogenesis pathway and are essential for spermatogenesis in mice, we asked whether the PIWI proteins Siwi and BmAgo3 might also be involved in the regulation of spermatogenesis in *B. mori*. To explore this possibility, we first performed qRT-PCR analysis of *Siwi, BmAgo3* and *BmHen1* using the samples prepared from WT sperm bundles grown till the fourth day of the fifth instar. We found that *Siwi, BmAgo3*, and *BmHen1* were all expressed in sperm bundles, with *BmHen1* being expressed 2-4 times higher than *Siwi* and *BmAgo3* (Fig 4A). We then constructed the loss-of-function mutants of *Siwi* and *BmAgo3* (*ΔSiwi* and *ΔBmAgo3*) by using a binary CRISPR/Cas9 system. To this end, we designed two sgRNAs targeting exons 1 and 3 of *Siwi* and *BmAgo3*, and *ΔSiwi* and *ΔBmAgo3* mutants were generated through genetic crossing between the

**Fig 4.**
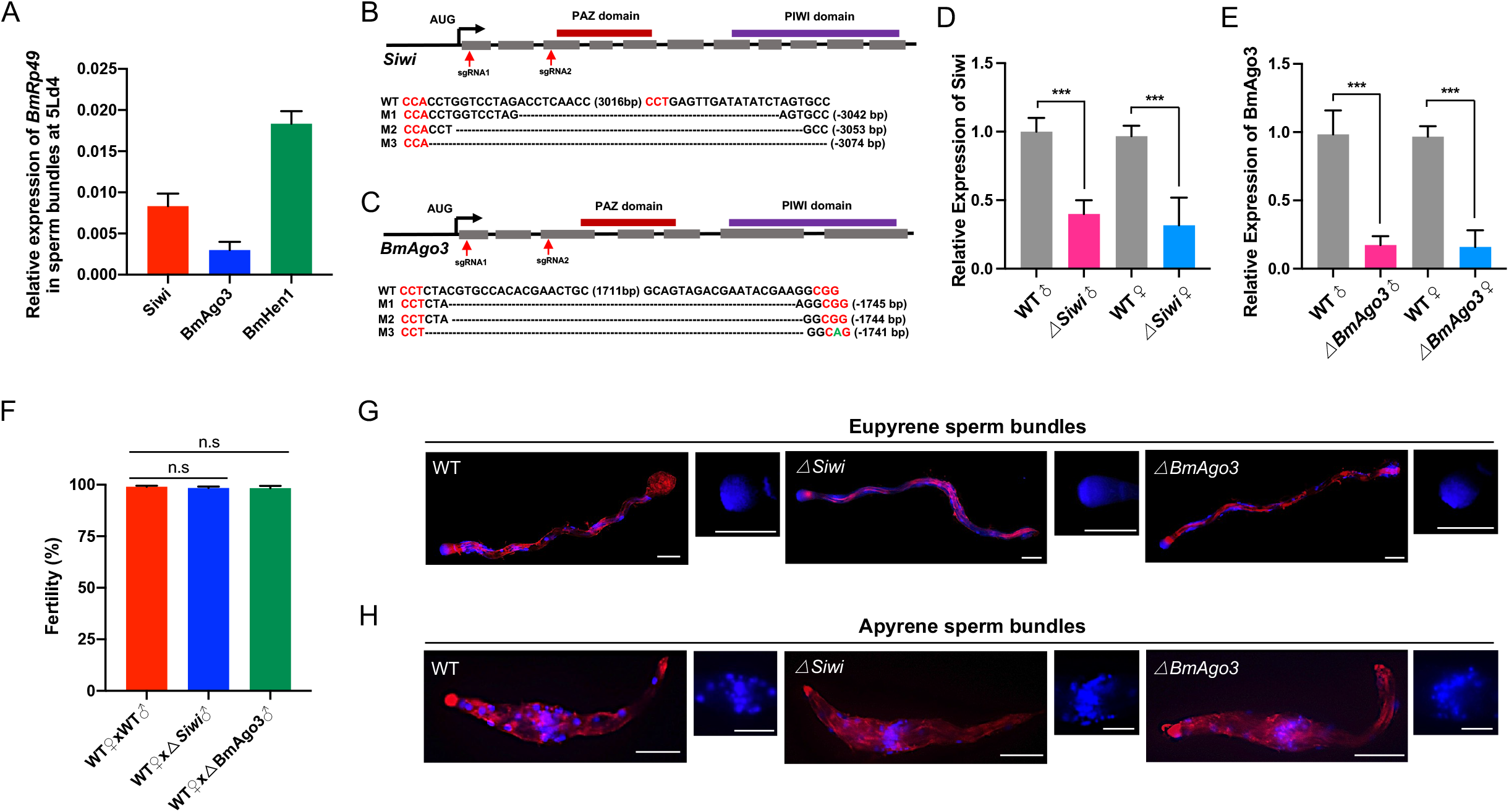
The PIWI proteins Siwi and BmAgo3 are dispensable for spermatogenesis in *Bombyx mori*. (A) Transcript levels of *Siwi, BmAgo3*, and *BmHen1* on day four of the fifth instar. (B and C) Genomic disruption of *Siwi* and *BmAgo3* gene using CRISPR/Cas9. Schematic of *Siwi* (B) and *BmAgo3* (C) genes structure and sgRNA targets. The exons of ORFs are represented by gray filled boxes, and the function domains (PAZ domain, PIWI domain) predicted by SMART are highlighted with red and purple boxes above. Red arrows indicate the target sites of sgRNA1 and sgRNA2. Genomic mutations of the *Siwi* and *BmAgo3* genes are shown with target sequences. The dashed lines indicate deleted sequences. M1-M3 represents three independent mutant lines. (D) and (E) qRT-PCR analysis of the transcript levels of *Siwi* and *BmAgo3* detected in testis and ovary of WT, *ΔSiwi*, and *ΔBmAgo3* at the wandering stage, respectively. Three individual biological replicates were performed, and error bars are mean ± SEM. The asterisks (***) indicate significant differences relative to WT (*P* < 0.001, t-test). (F) Fertility of males and females of the indicated genotypes. Fertility is indicated on the histogram (n= 15, *P* < 0.001, Fisher exact test). (G and H) Representative immunostaining images of eupyrene sperm bundles (G) and apyrene sperm bundles (H) from WT, *ΔSiwi*, and *ΔBmAgo3* on the seventh day at the pupal stage. Scale bars, 50 μm.

U6-sgRNA lines and the nos-Cas9 lines, respectively (Fig 4B and C). Mutations in the randomly selected representative F1 offspring were detected by genomic PCR and DNA sequencing, which indicates that the ORFs of *Siwi and BmAgo3* were shifted in both female and male *ΔSiwi* and *ΔBmAgo3* individuals, respectively (Fig 4B and C). Further qRT-PCR analysis showed that *Siwi* and *BmAgo3* were hardly expressed in *ΔSiwi* and *ΔBmAgo3*, respectively (Fig 4D and E), confirming that they are loss-of-function mutants.

Consistent with our previous report (Li et al., 2018), the *ΔSiwi* and *ΔBmAgo3* females produced much fewer eggs than WT. We then generated the hybrids by crossing *ΔSiwi* and *ΔBmAgo3* male with WT female, respectively, and performed fecundity tests. The results showed that they all were male-fertile (Fig 4F). Further fluorescence staining assay confirmed that *ΔSiwi* and *ΔBmAgo3* mutants had no defects in both eupyrene and apyrene sperm bundles (Fig 5G and H). Taken together, these results demonstrate that Siwi and BmAgo3 are not required for spermatogenesis in *B. mori*.

**Fig 5.**
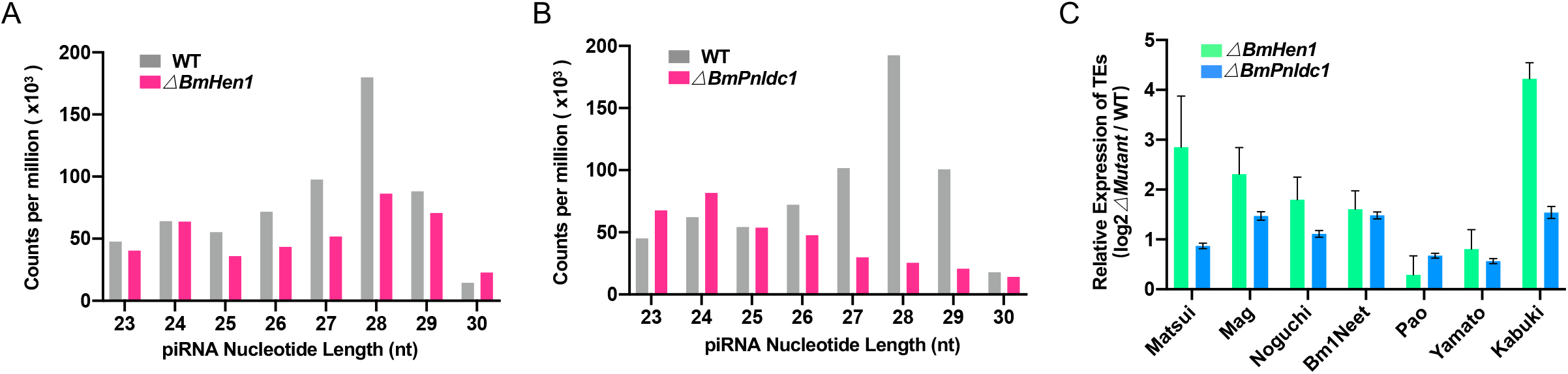
BmHen1 and BmPnldc1 are essential for piRNA biogenesis and suppression of transposon activity in sperm bundles. (A and B) Size distribution of piRNAs in *ΔBmHen1* (A) and *ΔBmPnldc1* (B) sperm bundles compared to WT. (C) Fold change of transposons in *ΔBmHen1* and *ΔBmPnldc1* sperm bundles compared to WT. TEs, transposon elements.

### 3.5 BmHen1 and BmPnldc1 are essential for piRNA biogenesis and suppression of transposon activity in sperm bundles

It is shown that, only after piRNA is trimmed by BmPnldc1, BmHen1 is able to modify 2’-O-methylation at its 3’ end (Izumi et al., 2016; Kawaoka et al., 2011). Knockdown of either *BmHen1* or *BmPnldc1* reduces 2’-O-methylated piRNAs and induces their degradation. These studies indicate that BmHen1 and BmPnldc1 act in the same pathway to modulate and stabilize piRNA (Izumi et al., 2016). To investigate whether BmHen1 and BmPnldc1 might participate in piRNA biogenesis during sperm development, we first performed small RNA-sequencing (sRNA-seq) assays using spermatocytes and sperm bundles isolated from testis of WT, *ΔBmHen1* and *ΔBmPnldc1* mutants, respectively. piRNA length distribution analysis showed that the abundance of 26-29 nucleotide long piRNA was remarkably reduced in*ΔBmHen1* and *ΔBmPnldc1* mutants (Fig 5A and B), indicating a critical role for both BmHen1 and BmPnldc1 in the regulation of piRNA stability in sperm bundles.

Next, we evaluated the effects of BmHen1 and BmPnldc1 on transposon activity using *ΔBmHen1* and *ΔBmPnldc1* mutants, and found that the activity of lots of transposons was depressed in the sperm bundles of these mutants (Fig 5C). We further examined the mRNA levels of seven well-annotated transposons by qRT-PCR. The results showed that 5 and 7 transposons were up-regulated in *ΔBmHen1* and *ΔBmPnldc1* mutants, respectively (Fig 5D). Altogether, these data suggest that both BmHen1 and BmPnldc1 are involved in stabilizing piRNA and activating transposon activity in sperm bundles in *B. mori*.

### 3.6 BmHen1 and BmPnldc1 co-regulate many genes in the same direction

As *ΔBmHen1* and *ΔBmPnldc1* mutants show similar defects in sperm development (Chen et al., 2020), we asked whether they might co-regulate the same set of downstream genes. To test this possibility, we performed RNA-sequencing (RNA-seq) assays using *ΔBmHen1* and *ΔBmPnldc1* mutants to gain potential insights into the changes in the expression of global genes associated with the defects in male fertility observed for *ΔBmHen1* and *ΔBmPnldc1* mutants. The results showed that there are up to 821 and 326 differentially expressed genes (DEGs) regulated by BmHen1 and BmPnldc1, respectively (Table 2 and 3). Among them, 157 DEGs are co-regulated by BmHen1 and BmPnldc1, of which 124 (78%) and 8 (19%) are up-regulated and down-regulated by BmHen1 and BmPnldc1, respectively (Fig 5A). Next, we performed gene ontology (GO) enrichment analysis, and found that the genes regulated by BmHen1 and BmPnldc1 are preferentially associated with cellular structure pathways including dynein complex, ciliary related, and membrane components (Fig 5B). We filtered the unannotated and transposon-related DEGs co-regulated by BmHen1 and BmPnldc1, and then generated a heat map through hierarchical clustering analyses, which reveals that 17 of these genes are regulated by BmHen1 and BmPnldc1 in the same direction (Fig 5C). Among them, the genes participating in energy metabolism and cell differentiation are significantly up-regulated in both *ΔBmHen1* and *ΔBmPnldc1* mutants, which include *glutamate receptor ionotropic* (*GluR*), *UDP glycosyltransferase precursor* (*UGT*), *ATP synthase mitochondrial complex* (*ATPase*), *intraflagellar transport protein* (*IFT*) and *cyclin-dependent kinase* (*CDK*). We then verified the expression of *GluR, UGT, ATPase, IFT*, and *CDK* in WT, *ΔBmHen1*, and *ΔBmPnldc1* mutants through qRT-PCR, respectively, and found that all these genes expression was dramatically enhanced in *ΔBmHen1* and *ΔBmPnldc1* mutants (Fig 6D and E). These results, in conjunction with those from the phenotype analysis, indicate that BmHen1 and BmPnldc1 regulation of eupyrene spermatogenesis may be achieved through modulation of the genes involved in energy metabolism and cell differentiation.

**Fig 6.**
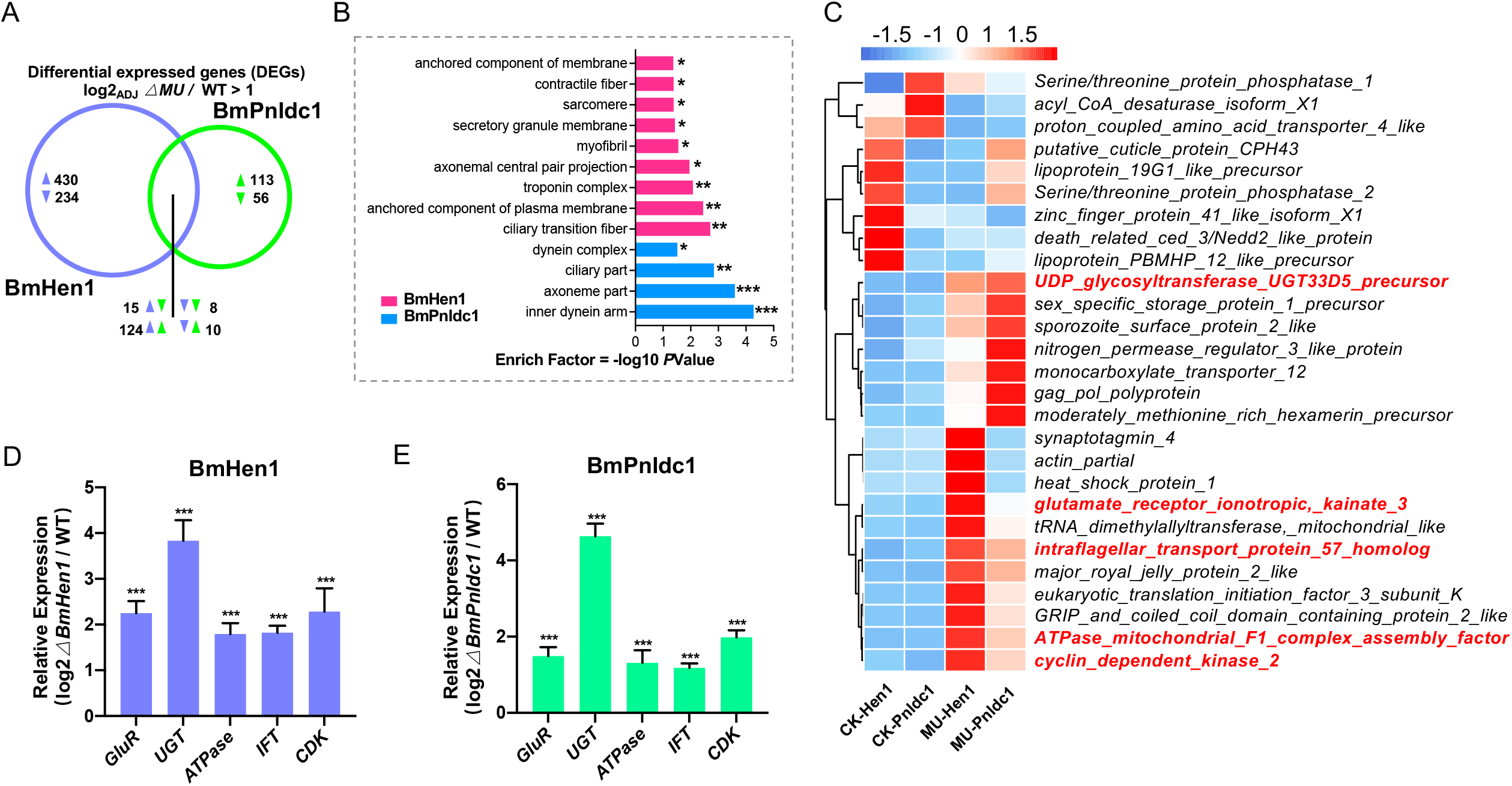
BmHen1 and BmPnldc1 co-regulate many genes in the same direction. (A) Venn diagram analysis showing DEGs in sperm bundles commonly and uniquely regulated by BmHen1 (left, orange) and BmPnldc1 (right, green). (B) Gene ontology enrichment analysis showing the biological processes most associated with the DEGs regulated by BmHen1 and BmPnldc1. Enrichment analysis was conducted using the functional annotation tool DAVID. Statistical significance was determined at a p-value ≤ 0.05, *; p-value ≤ 0.01, **; p-value ≤ 0.001, ***; (C) Hierarchical clustering of selected spermatogenesis-related DEGs in different samples. CK-Hen1, control of BmHen1; CK-Pnldc1, control of BmPnldc1; MU-Hen1, *ΔBmHen1* mutant; MU-Pnldc1, *ΔBmPnldc1* mutant. (D and E) qRT-PCR analysis of the selected DEGs regulated by BmHen1 (D) and BmPnldc1 (E). Asterisks (***) indicate *P* < 0.001 determined by a two-tailed Student’s t-test. Data are mean ± SEM. https://doi.org/10.1126/science.1107130

## Discussion

It is known that the piRNA biogenesis pathway is involved in germline development in the animal kingdom (Juliano et al., 2011; Ozata et al., 2019). Loss-of-function of the components in piRNA pathway typically leads to oogenesis defects and female sterility in *Drosophila* or male gonad development defects in mice (Juliano et al., 2011) (Juliano et al., 2011). In our previous studies, we have genetically linked genes involved in piRNA biogenesis such as *BmMael* and *BmPnldc1* to sperm development in *B. mori*, and demonstrated that BmMael and BmPnldc1 are involved in regulating spermatogenesis in *B. mori* (Chen et al., 2019; S. Chen et al., 2020). We were interested in exploring whether other key components in the piRNA pathway such as BmHen1 and PIWI proteins Siwi and BmAgo3 might play a role in the regulation of sperm development. In the current study, we identified BmHen1 as a new important component involved in the regulation of eupyrene sperm development in *B. mori*. We generated the loss-of-function mutants of BmHen1 through a binary transgenic CRISPR/Cas9 system (Fig 1B). Phenotype analyses of *ΔBmHen1* mutant demonstrated that it is required for both female and male fertility (Fig 2D). Further fluorescence staining assays demonstrated that the male sterility is attributed to eupyrene sperm defects occurring during the elongation stage (Fig 3A and C). These results demonstrate that BmHen1 plays an essential role in the regulation of eupyrene sperm development.

In our previous study, we have characterized a role for PIWI proteins Siwi and BmAgo3 in regulating oogenesis. (Li et al., 2018). In the present study, we explored whether Siwi and BmAgo3 might regulate eupyrene sperm development in in *B. mori*. Unexpectedly, Siwi and BmAgo3 are not required for spermatogenesis in *B. mori*. The evidence supporting this conclusion comes from our genetic studies, in which the loss-of-function mutant of either *Siwi* or *BmAgo3* is male-fertile (Fig 4F), and fluorescence staining assays, in which both morphs of sperms are normal in either *ΔSiwi* or *ΔBmAgo3* mutant (Fig 4G and H). As the core elements of the piRNA pathway, PIWIs are loaded with piRNAs and play a critical role in both piRNA biogenesis and piRNA-induced gene silencing (Czech et al., 2018; Iwasaki et al., 2015; Ozata et al., 2019). During piRNA biogenesis, Hen1 modifies a 2’-O-methylation at the 3’ end of piRNAs, which inhibits their degradation (Horwich et al., 2007; Kamminga et al., 2010; Kirino and Mourelatos, 2007; Saito et al., 2007). Why BmHen1 is essential for eupyrene sperm development in *B. mori* but the PIWI proteins Siwi and BmAgo3 are not awaits further investigation in future studies.

How may BmHen1 participates in eupyrene sperm development in *B. mori*? It is known that, in *Drosophila*, Hen1 not only acts to modify piRNAs but also siRNAs. However, it is reported that the 3’ end of siRNAs in lepidopterans lacks the 2’-O methylation (Fu et al., 2018). Perhaps BmHen1 is involved in the modification of an unidentified class of small RNAs, which is crucial for sperm development in *B. mori*. In recent years, a new class of small RNAs so-called tRNA-derived interference RNAs (tsRNAs) has been reported to be important for sperm development in mouse (Chen et al., 2016b). Dysregulation of tsRNAs can affect sperm bioactivity and lead to male sterility (Chen et al., 2016a). Sperm tsRNAs are mainly derived from the 5’ end of tRNAs, ranging from 29 nt to 34 nt in size, which are more abundant than miRNAs constituting the majority of small RNAs in sperm (Kim et al., 2017). Although the biogenesis mechanism of tsRNAs is intricate and still unclear, the genes encoding endonuclease and methyltransferase have been linked to this process (Pan et al., 2021). In mouse spermatogonial stem cells, some tRNA-derived small RNAs are known to undergo Hen1-dependent methylation (Peng et al., 2018). It will be interesting to investigate whether BmHen1 may regulate sperm development through modulation of tsRNA biogenesis in furture studies.

In our study, we observed that the defects in *ΔBmHen1* eupyrene sperm are similar to those in*ΔBmPnldc1* eupyrene sperm (Fig 3A and C), which indicates an equal essential role for BmHen1 and BmPnldc1 in regulating eupyrene sperm development in *B. mori*. BmPnldc1 was initially identified in C elegans and BmN cells, which encodes an exonuclease, trimming piRNAs at their 3’ end (Izumi et al., 2016; Tang et al., 2016). It is reported that the 2’-O-methylation caused by Hen1 occurs in a manner coupled with trimming which indicates the biochemical function of BmHen1 depends on BmPnldc1 (Izumi et al., 2016). We also demonstrated that both BmHen1 and BmPnldc1 are essential for piRNA biogenesis and suppression of transposable elements in sperm bundles (Fig 5A and B), which is consistent with previous studies performed in BmN cells. Further RNA-seq analysis suggests that BmHen1 and BmPnldc1 regulation of eupyrene sperm development may be achieved, at least in part, through modulation of the genes involved in energy metabolism and cellular differentiation (Fig 6B and C). Given our hypothesis that BmHen1 may act in tsRNAs processing, it is possible that BmPnldc1 is also involved in this process. For example, BmPnldc1 acts as an exonuclease to trim tsRNAs precursors into mature tsRNAs, and then BmHen1 catalyzes a 2’-O-methylation at the 3’ end of the mature tsRNAs to inhibit their degradation. As a result, the mature tsRNAs accumulate, and may regulate the expression of genes such as *GluR, UGT, ATPase, IFT*, and *CDK*, and eventual spermatogenesis in *B. mori*. It will be worth exploring these possibilities in further studies.

## Acknowledgments

We thank Prof. Hong-Quan Yang for proofreading the manuscript. This research was supported by the National Science Foundation of China (32021001) and Strategic Priority Research Program of Chinese Academy of Sciences (Grant No. XDPB16).

